# Atomic Force Microscopy reveals differences in mechanical properties linked to cortical structure in mouse and human oocytes

**DOI:** 10.1101/2025.01.14.632898

**Authors:** Rose Bulteau, Lucie Barbier, Guillaume Lamour, Yassir Lemseffer, Marie-Hélène Verlhac, Nicolas Tessandier, Elsa Labrune, Martin Lenz, Marie-Emilie Terret, Clément Campillo

**Affiliations:** LAMBE, Univ Evry, CNRS, Université Paris‐Saclay, 91025, Évry-Courcouronnes, France; Center for Interdisciplinary Research in Biology (CIRB), Collège de France, Université PSL, CNRS, INSERM, 75005 Paris, France; LPTMS, CNRS, Université Paris-Sud, Université Paris-Saclay, 91405 Orsay, France; Hospices Civils de Lyon, service de médecine de la reproduction et préservation de fertilité; Inserm U1208, SBRI, Université Claude Bernard Lyon 1, faculté de médecine Laennec, France; Institut Universitaire de France (IUF), 75005 Paris, France

**Author notes:** These authors contributed equally to this work (Rose Bulteau, Lucie Barbier).

**Keywords:** Atomic Force Microscopy, Actomyosin cortex, Elastic modulus, Cortical tension, Oocytes, Biomechanics

## Abstract

Cell mechanical properties regulate biological processes such as oocyte development. Cortical tension is regulated via actomyosin cortex remodeling to ensure optimal oocyte quality. However, the evolution of other mechanical parameters and their relationship with cortex structure remain poorly understood in mammalian oocytes. In this work, we propose a methodology combining multiple mechanical parameters measured through Atomic Force Microscopy to investigate the relationship between oocyte mechanical properties and cortex organization. By studying mouse oocytes at various stages of development, along with engineered ones with specific cortex organization, we demonstrate that a thin actin cortex corresponds to stiff oocytes while a thick one is associated with softer oocytes. We further reveal that maternal age, a critical factor for fertility, affects mouse oocytes mechanics correlating with alterations in their cortex structure. Finally, we show that the evolution of mechanical properties differs between human and mouse oocyte development, highlighting species-specific differences in cortex organization.

## INTRODUCTION

Living cells are complex materials whose mechanical properties evolve during biological processes such as cell division^[1]^ and during the evolution of pathologies such as cancer^[2,3]^. Mechanical properties of the cell surface are modulated by the cell cortex, an actin-rich structure lying beneath the plasma membrane, in which actin filaments are nucleated at the membrane and interact with myosin motors^[4]^. Cortical tension is a mechanical property that corresponds to the energy required to increase the cell surface^[5]^ and is determined by the dynamics of actin polymerization and myosin activity^[6]^. It has been shown that cortical tension is tightly regulated during the final phase of mouse oocyte development, called meiotic maturation^[7,8]^. At birth in mammals, oocytes are arrested in prophase of the first meiotic division (prophase I, PI). After puberty and cyclically, some oocytes undergo meiotic maturation: they exit from prophase I, complete the first meiotic division (meiosis I, MI), and arrest in metaphase of the second meiotic division (meiosis II, MII) until fertilization. After meiosis I entry, the oocyte cortex gradually thickens through actin nucleation, chasing cortical myosin-II and leading to a decrease in cortical tension^[7]^. Aberrant low or high cortical tension, a frequent defect in mammalian oocytes^[9]^, can hinder division geometry and induce aneuploidy^[10,7,11]^. These studies show that the actomyosin cortex is a key factor for proper oocyte formation and that cortical tension measurements may indicate alterations in the actomyosin cortex structure. However, how other mechanical parameters, such as elasticity and viscosity, evolve with changes in the actomyosin cortex is still unknown.

Atomic Force Microscopy (AFM) is a robust method for probing multiple mechanical properties of living cells surface^[3,12–20]^. In AFM experiments, the cell is probed by a nanometer-sized tip located at the end of a flexible cantilever. To measure mechanical parameters, the tip presses on the cell, inducing a local deformation. The force applied and the resulting cell deformation are recorded through the cantilever deflexion, producing force-indentation curves. Fitting these force-indentation curves with appropriate physical models gives access to the cell’s mechanical properties. In most studies, living cells are treated as bulk elastic materials, and their effective elastic modulus is extracted from these curves^[21,22]^. Only a few studies use local AFM indentation to measure cell cortical tension^[23,24]^. The respective contributions of elastic modulus and tension on the cell cortex response in AFM experiments are unclear. Moreover, the cell internal medium also dissipates energy, as shown in AFM experiments by the difference between approach and retract curves^[25]^. Few works have assessed oocyte mechanical properties using AFM. In most cases, they have probed the *Zona Pellucida*, the porous network of glycoproteins constituting the extracellular matrix surrounding mammalian oocytes, using purely elastic models^[26–28]^. Some have probed the mechanical contributions associated with the *Zona Pellucida* and the whole oocyte^[29,30]^, using time-relaxation measurements with flat macro-cantilevers^[28]^. However, AFM measurements directly probing the oocyte cortex have not been performed and could extend the current description of oocyte mechanics. Moreover, existing mechanical studies of oocytes are limited to a single mechanical parameter, such as elastic modulus or cortical tension.

Here, we propose a methodology that combines several mechanical parameters obtained through Atomic Force Microscopy experiments to study the links between oocyte’s mechanical properties and actin cortical organization. We developed an elasto-capillary model and analysis pipeline that allows extracting oocyte elastic modulus, cortical tension, and energy dissipation. We confirmed that our model accurately describes oocyte force-indentation response and yields values of cortical tension consistent with the ones obtained by micropipette aspiration^[7,8,31]^. We investigated different stages of mouse oocyte maturation and engineered mouse oocytes in which the actomyosin cortex is manipulated to obtain oocyte models with increased or decreased cortical tension. Using principal component analysis combining the parameters obtained by AFM, we found that groups of oocytes cluster depending on their actin cortical organization. Importantly, we also describe this link between actin cortical organization and mechanical properties in the subfertile case of mouse advanced maternal age and in human oocytes. However, the evolution of the mechanical properties during meiotic maturation differs between human and mouse oocytes, suggesting different molecular regulation.

## RESULTS

### Describing oocyte mechanics with an elasto-capillary model

We used the AFM measurement technique adapted to mammalian oocytes, which we recently developed^[32]^ (Figure 1A). To probe the oocyte cortex without the mechanical contributions associated with the *Zona Pellucida*, we performed the measurements after removing this layer (Figure S1A). We indented the top of the oocyte and monitored the compression force ℱ as a function of the oocyte indentation *δ*_tot_ We first gradually compressed the oocyte by lowering the indenter over the course of seconds up to a maximal applied force set at 0.5 nN, then raised the indenter again. We thus collected two force-indentation curves corresponding to the approach and the retract phases (Figure 1B). From these force curves, we can extract several mechanical parameters.

**Figure 1.**
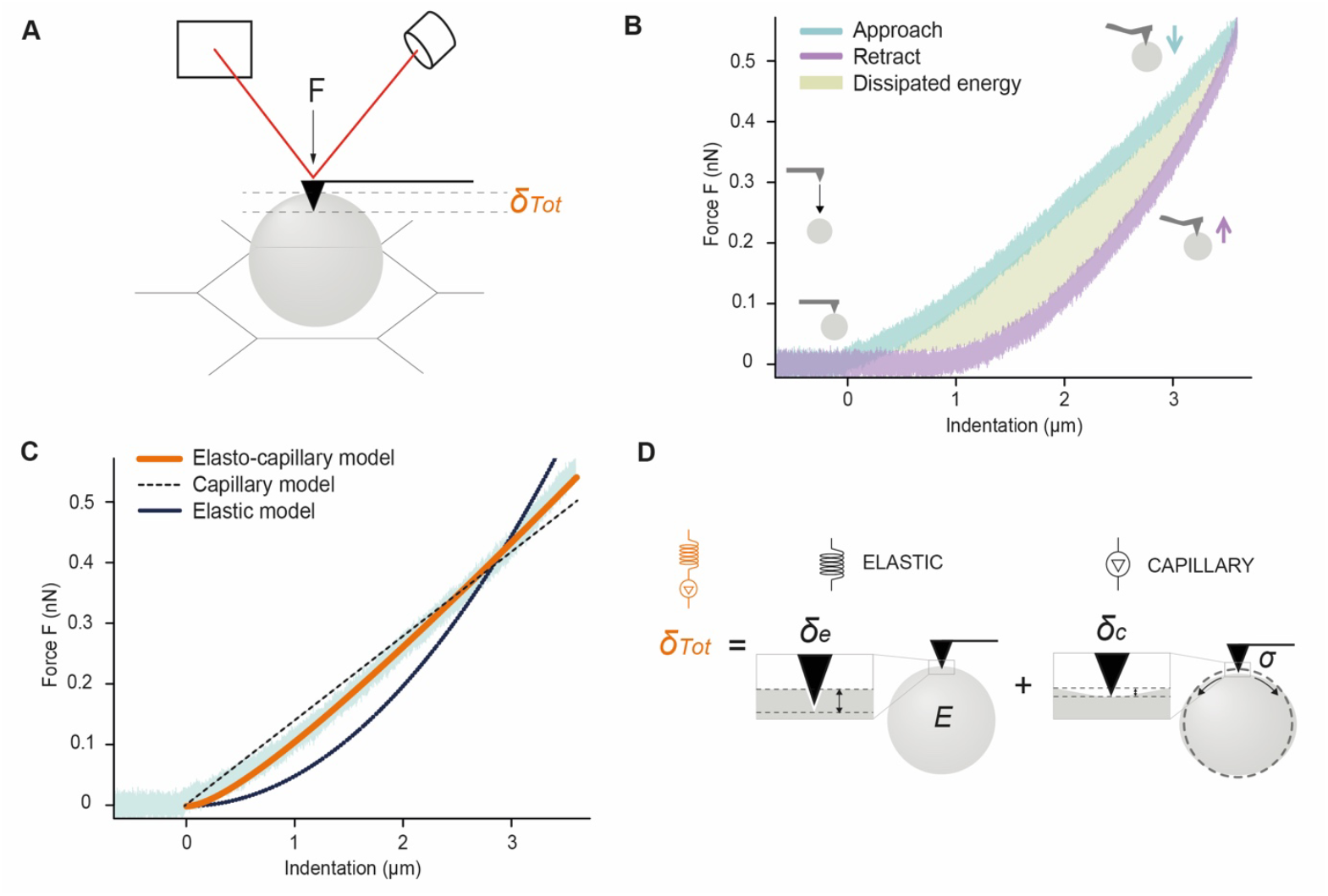
AFM measurement of mammalian oocyte mechanics: strategy, modeling, and analysis. **A.** Scheme of the AFM measurement technique (oocyte in grey placed on an electronic microscopy grid, laser in red). **B.** Example of a representative approach (blue) and retract (purple) force-indentation curve obtained for one oocyte. The green area between the approach and retract curves corresponds to the dissipated viscous energy. **C.** Graph showing the force-indentation curve fits with an elastic model (continuous black line), a capillary model (black dashed line), and the elasto-capillary model (orange line), which is used to extract oocyte cortical tension and elastic modulus. **D.** The total indentation in the oocyte (∂_*Tot*_) is the sum of the elastic (∂_*e*_) and capillary (∂_*c*_) indentations. The oocyte (in grey) is described mechanically as a spring and a contractile element in series.

The approach and retract curves do not overlap, revealing a viscous energy dissipation during the indentation. We calculate this dissipated energy by integrating the area between the approach and the retract curves (Figure 1B). This quantity is normalized by the area under the approach curve^[25]^. It thus provides a dimensionless quantity close to 1 in cases of high viscous dissipation and 0 for low dissipation. These values range from 0.4 to 0.6, showing that both dissipative and non-dissipative contributions must be considered in the oocyte’s mechanical description.

To infer information about the non-dissipative part of the oocyte’s mechanical response, we consider the approach indentation curve and note that it combines characteristics of the response of two distinct non-dissipative elements, one at small indentation and the other at large ones (Figure 1C). For small indentation, it displays a parabolic dependence of the force on the indentation. This is consistent with the response of an elastic homogeneous and isotropic half-space indented by a pyramidal probe. In such a setting, the standard Hertz model predicts that the indentation force depends on the indentation *δ*_*e*_ of the elastic half-space through:

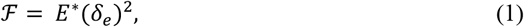

where the characteristic modulus 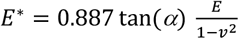 is a function of the elastic modulus *E*; the Poisson ratio *ν* = 0.5 (assuming an incompressible material) and the side angle α = 17.5° of the indenter cone. For larger indentations, the force appears to depend affinely on the indentation. This is consistent with the capillary response of a disk of membrane-like thin elastic slab indented over a depth *δ*_*c*_, namely:

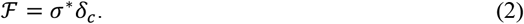

This expression involves a characteristic tension 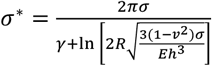, which features the cortical tension *σ*, the Euler gamma constant *γ* ≃ 0.577, the radius *R* of the membrane disk and the thickness *h* of the membrane. While *σ*^*^ depends on many parameters, for large enough membranes the logarithmic dependence of its denominator is very weak and thus *σ*^*^, for typical values of experimental parameters (*R* = 40*μm, h* = 2*μm*, E=1kPa, *σ* =1nN.μm^-1^), is equal to the cortical tension *σ*.

We rationalize this complex dependence of the force on the indentation through a geometric argument. We assimilate the oocyte cortex to a tense elastic shell whose thickness *h* is much smaller than its radius *R*. In such a setting, the cortex deformation induced by the indenter involves two widely different length scale (Figure 1D). At a very local scale, the tip of the indenter penetrates into the cortex by a distance *δ*_*e*_, *i*.*e*., the distance between the tip and the cortex mid-plane is *h*/2 − *δ*_*e*_. On larger length scales, the downward force exerted by the indenter additionally pushes this midplane down by a distance *δ*_*c*_, implying that over length scales much larger than *h* the cortex acts as a tense membrane subjected to a downward point force. We reason that the total indentation *δ*_tot_ of the oocyte is the sum of these local and large-scale displacements, implying

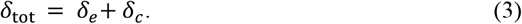

Inserting Eqs. (1-2) yields the following expression for the total indentation:

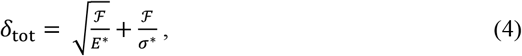

which leads to:

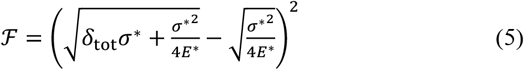

with which we fit our approach curve (Figure 1C, 1D). Eq. (3) is parabolic at low indentation and thus dominated by the elastic response. As expected, this expression crosses over from a parabolic elastic-dominated response for 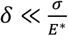 to a linear capillary regime for 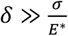. This elasto-capillary model accurately fits our force curves in the approach phase (Figure 1D), and the Residual Standard Errors (RSE) is lower as compared to elastic or capillary models (Figure S1B). Therefore, our measurements simultaneously provide values of oocyte elastic modulus *E* and cortical tension *σ*.

To quantify the elastic vs. capillary character of the oocytes in our conditions of indentation, we introduce a dimensionless parameter, which we name the capillary indentation ratio *r*_*c*_:

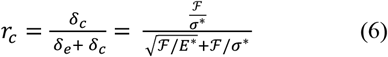

A ratio *r*_*c*_ = 0 indicates that the oocyte has a purely elastic behavior. In contrast, *r*_*c*_ = 1 corresponds to a purely capillary response. For 70% of our data, 0.05 < *r*_*c*_ < 0.95 at the maximum indentation force, implying a mixed behavior. In other few cases, one of the two behaviors dominates the oocytes’ mechanical response, implying that their response curves contain very little information about the other. In practice, in such cases, our fitting procedure returns unreliable values for the elastic parameter associated with the subdominant behavior (Figure S1C). We thus disregarded these curves in the following.

Our AFM experiments and fitting procedure thus provide four mechanical parameters characterizing oocytes: normalized dissipated energy (*DE*), cortical tension (σ), elastic modulus (*E*), and maximum capillary indentation ratio (*r*_*c*_).

### Oocyte mechanics predict cortex organization

To validate our model and investigate the link between oocyte mechanics and cortex organization, we first probed mouse oocytes with known cortical tension and organization. We compared oocytes arrested in prophase I (PI), oocytes in meiosis I (in early or late MI), oocytes arrested in meiosis II (MII), and engineered oocytes with a modified cortex organization^[7]^ (Figure 2A). These oocytes express cortical probes either forcing cortical actin nucleation and chasing cortical myosin-II (cVCA, lower cortical tension^[11]^ or forcing myosin-II recruitment at the cortex (cRhoA^[34]^, higher cortical tension - Figure S1D). As a control, we used oocytes expressing only the cortical anchor (Ctrl). We characterized the mechanics of all these types of oocytes using AFM.

**Figure 2.**
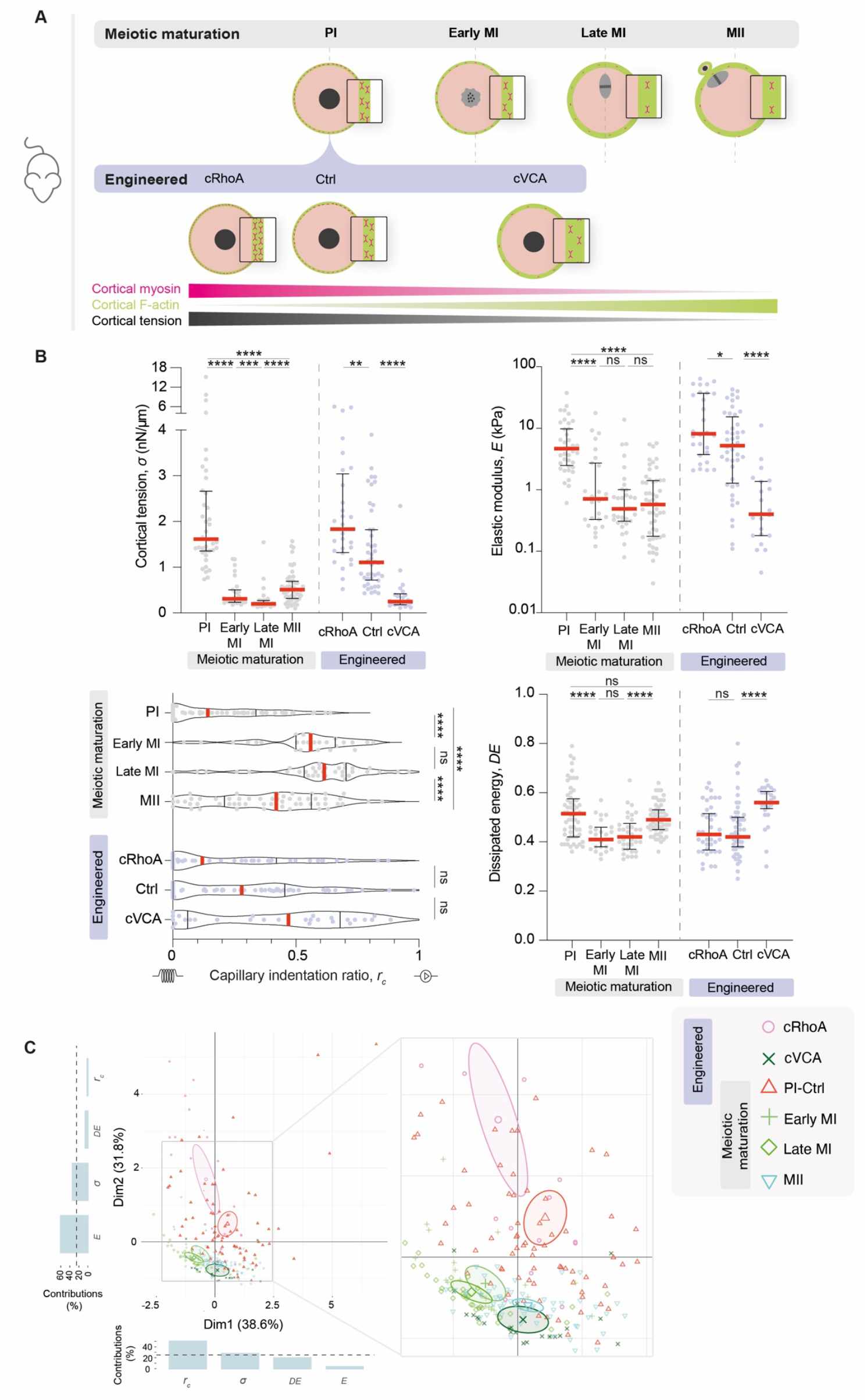
Oocyte mechanics predict cortex organization. **A.** Scheme of the different oocytes with known cortical organization and known cortical tension measured with AFM, displaying cortex organization (cortical actin in green, cortical myosin-II in pink). The upper row shows oocytes at different stages of meiotic maturation (prophase I: PI, early meiosis I: early MI, late meiosis I: late MI, meiosis II: MII). The middle row shows engineered oocytes in prophase I with a modified cortex (cVCA, cRhoA), with their control (Ctrl) expressing only a cortical anchor. The lower row shows the enrichment of actin and myosin-II in the cortex of the oocytes displayed above, along with their cortical tension (measured previously with micropipette aspiration). The nucleus is shown as a black sphere, the microtubule spindle as a gray oval, and the chromosomes as black spheres. **B.** Evolution of mechanical parameters during meiotic maturation (PI, early MI, late MI, MII) and for engineered oocytes (cRhoA, cVCA, Ctrl) in prophase I. Graphs showing cortical tension α (nN.μm^-1^), elastic modulus E (kPa), capillary indentation ratio r^c^ (a.u), dissipated viscous energy DE (a.u). Data are medians and quartiles with individual data points plotted from at least three experiments. For cortical tension and elastic modulus, 40 PI, 27 early MI, 33 late MI, 53 MII, 28 cRhoA, 44 Ctrl, and 22 cVCA were measured. For capillary indentation ratio, 58 PI, 27 early MI, 34 late MI, 57 MII, 42 cRhoA, 58 Ctrl, and 29 cVCA were measured. For the normalized dissipated viscous energy, 58 PI, 27 early MI, 34 late MI, 57 MII, 42 cRhoA, 57 Ctrl and 29 cVCA were measured. The statistical significance between the two groups was assessed with an unpaired t-test or a nonparametric Mann-Whitney test, depending on whether the data followed a Gaussian distribution. **C.** Principal component analysis of the four mechanical parameters for the different stages of meiotic maturation and engineered oocytes (20 cRhoA, 77 PI-Ctrl, 22 cVCA, 27 early MI, 33 late MI and 53 MII). The graphs show the percentage contributions of the four mechanical parameters in PCA Dimension 1 (Dim-1) and Dimension 2 (Dim-2). The black dotted line is 25%.

We first measured a decrease in cortical tension after meiosis I entry (Figure 2B, 1.6 nN.μm^-1^ for PI vs. 0.2 nN.μm^-1^ for late MI) and for cVCA oocytes (Figure 2B, 0.25 nN.μm^-1^ for cVCA vs. 1.1 nN.μm^-1^ for Ctrl). By contrast, we measured an increase in cortical tension for cRhoA compared to Ctrl oocytes (Figure 2B, 1.8 nN.μm^-1^ for cRhoA vs. 1.1 nN.μm^-1^ for Ctrl). These results are consistent with previous measurements of cortical tension obtained with micropipette aspiration^[7,8,10,11]^, validating the elasto-capillary model.

Then, we assessed the three additional mechanical parameters described above. After meiosis I entry, the elastic modulus decreases progressively (4.6 kPa for PI vs. 0.5 kPa for late MI) while the capillary indentation ratio increases. Normalized dissipated energy also decreases after meiosis I entry but returns in Meiosis II to values similar to those of oocytes in prophase I (Figure 2B). For engineered oocytes, the elastic modulus is higher for cRhoA (8.2 kPa) and lower for cVCA (0.4 kPa) than for Ctrl oocytes (5.2 kPa). The normalized dissipated energy is higher for cVCA oocytes and similar between cRhoA and Ctrl oocytes. The capillary indentation ratio for cVCA oocytes is closer to a capillary-dominated response than for cRhoA ones (Figure 2B). Overall, while the elastic modulus seems to follow similar trends than cortical tension, the normalized dissipated energy and the capillary indentation ratio evolve independently in the different types of oocytes probed.

To facilitate the visualization of our multiparametric data set and possibly identify groups of oocytes with similar mechanical properties, we combined the four mechanical parameters in a principal component analysis (PCA) (Figure 2C). For all PCA, each ellipse represents the confidence intervals of the centroid position of the oocyte population. Two groups stand out: the first one is composed of cRhoA oocytes and prophase I oocytes (PI-Ctrl), and the second one is composed of cVCA oocytes, meiosis I and II oocytes. The first group clusters oocytes with thin actin cortices enriched in myosin-II (Figure 2A) and is associated with high cortical tension, high elastic modulus, and low capillary indentation ratio. In contrast, the second group clusters oocytes with thick actin cortices with fewer myosin-II (Figure 2A) and is associated with low cortical tension, low elastic modulus, and high capillary indentation ratio.

Our results highlight the tight relationship between cortex organization and mechanics. They suggest that oocyte mechanics could allow to infer cortex organization.

### Maternal age impacts oocyte mechanics and is associated with a modified cortex organization

Maternal age is a well-known factor affecting the ability of an oocyte to develop into a proper embryo^[35–37]^, but its effect on oocyte mechanics is unknown. We compared oocytes in prophase I obtained from young and aged female mice (11 and 44-56 weeks, respectively). We found that cortical tension and elastic modulus decrease with age (1.6 nN.μm^-1^ and 9 kPa for Young vs. 1.0 nN.μm^-1^ and 3 kPa for Aged, Figure 3A), whereas capillary indentation ratio increases and normalized dissipated energy is constant. In PCA, oocytes from aged mice are located in between the groups of oocytes with a thin actin cortex rich in myosin-II (cRhoA, PI-Ctrl and Young) and oocytes with a thick cortex and few myosin-II (meiosis I and cVCA) (Figure 3B).

**Figure 3.**
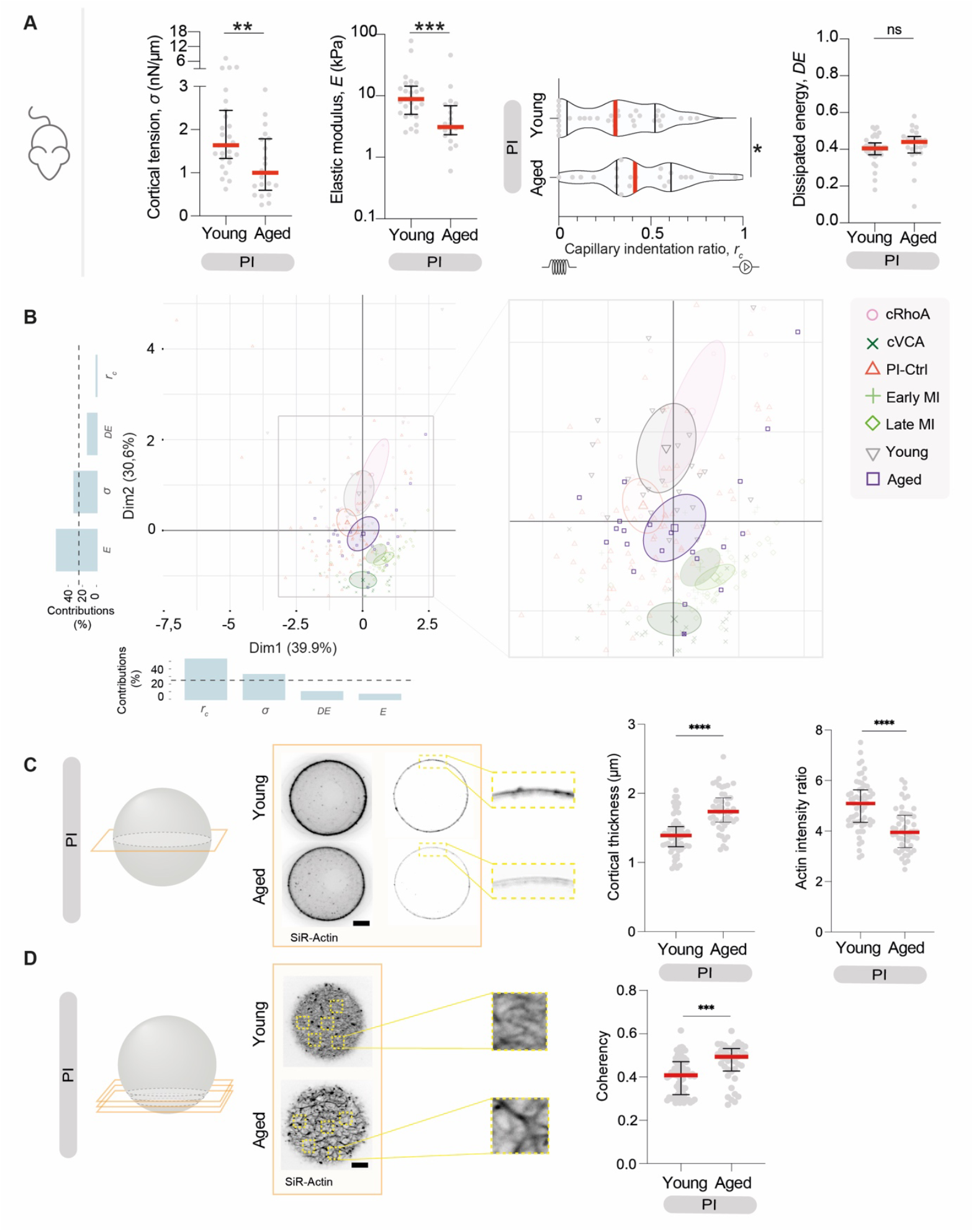
Maternal age impacts oocyte mechanics and is associated with a modified cortex organization. **A.** Graphs showing the mechanical parameters (cortical tension σ (nN.μm^-1^), elastic modulus E (kPa), capillary indentation ratio r^c^, normalized dissipated viscous energy DE (a.u)) for prophase I (PI) oocytes coming from Young (11-week-old) and Aged (44-56-week-old) mice. Data are medians and quartiles with individual data points plotted from three independent experiments. For cortical tension and elastic modulus, 27 oocytes from Young mice and 22 from Aged mice were analyzed. For capillary indentation ratio and normalized dissipated viscous energy, 34 oocytes from Young mice and 23 from Aged mice were analyzed. The statistical significance of differences between 2 groups was assessed with an unpaired t-test or a nonparametric Mann-Whitney test, depending on whether the data followed a Gaussian distribution. **B.** Principal component analysis of the four mechanical parameters for different stages of meiotic maturation, engineered oocytes, and oocytes coming from Young and Aged mice (20 cRhoA, 77 PI-Ctrl, 22 cVCA, 27 early MI, 33 late MI and 53 MII, 27 Young and 22 Aged). Ellipses indicate 95% confidence intervals. The graphs show the percentage contribution of the four mechanical parameters in PCA Dimension 1 (Dim-1) and Dimension 2 (Dim-2). The black dotted line is set at 25%. **C.** Confocal spinning disc images of prophase I (PI) oocytes coming from Young (11-week-old) and Aged (44-56-week-old) mice stained with SiR-Actin (black). Scale bar: 5 μm. One Z plane is shown. Graph showing the cortical thickness and the actin fluorescence ratio between the cortex and the cytoplasm of 47 prophase I (PI) oocytes coming from Young (11-week-old) and 59 prophase I (PI) oocytes coming from Aged (44-56-week-old) mice. Data are medians and quartiles with individual data points plotted from three independent experiments. The statistical significance of differences between 2 groups was assessed with a nonparametric Mann-Whitney test. **D.** Confocal spinning disc images of prophase I (PI) oocytes coming from Young (11-week-old) and Aged (44-56-week-old) mice stained with SiR-Actin (black). Scale bar: 10 μm. A projection of five Z planes is shown. Graph showing the coherency of cortex actin network calculated in 5.33 μm square ROIs for 67 prophase I (PI) oocytes coming from Young (11-week-old) and 40 prophase I (PI) oocytes coming from Aged (44-56-week-old) mice. Data are medians and quartiles with individual data points plotted from two independent experiments. The statistical significance of differences between 2 groups was assessed with an unpaired t-test.

We thus hypothesized that the cortex of oocytes in prophase I from aged mice resembles more that of cVCA or meiosis I oocytes. To test this hypothesis, we characterized the actin cortex of oocytes from young and aged mice. Indeed, oocytes from old mice present a thicker cortex than oocytes from young mice (Figure 3C), as for oocytes in late meiosis I (Figure S2A)^[7]^ or cVCA oocytes^[7,11]^. Furthermore, we found that cortical actin fluorescence intensity is lower in oocytes from aged mice (Figure 3C), suggesting a different actin organization. We confirmed this hypothesis by measuring a higher cortical coherency (see Materials and Methods) in these oocytes compared to those from young mice (Figure 3D, S2B).

Our results show that maternal age impacts oocyte mechanics: oocytes from aged mice are softer than those from young ones and harbor a differently organized and thicker actin cortex, further suggesting that oocyte mechanics reflect cortex organization.

### Human and mouse oocyte mechanics diverge, reflecting differences in cortex organization

We then adapted our AFM measurement and analysis pipeline to human oocytes, whose mechanics and cortex organization during meiotic maturation are unknown. In mouse oocytes, cortical actin thickening and tension decrease between prophase I and meiosis I are key events for successful oocyte division^[7,11]^. We therefore studied human oocytes in prophase I (PI), meiosis I (MI), and meiosis II (MII) to determine whether similar processes regulate their division.

The cortical tension of mouse and human oocytes is of a similar order of magnitude, albeit lower for human oocytes (Figure 4A). However, and contrary to mouse oocytes, cortical tension in human oocytes is similar in prophase I and meiosis I (Figure 4A, 0.6 nN.μm^-1^ for PI and MI), decreasing only in meiosis II (Figure 4A, 0.3 nN.μm^-1^ for MII). The elastic modulus of human oocytes does not exhibit significant changes across the three stages; the capillary indentation ratio increases in meiosis II, and the normalized dissipated energy increases progressively during meiotic maturation (Figure 4A). We performed a PCA including human and mouse oocytes (Figure 4B). Human oocytes in prophase I and meiosis I overlap and are distinct from human oocytes in meiosis II. These three ellipses are close to those of the cVCA mouse oocytes.

**Figure 4.**
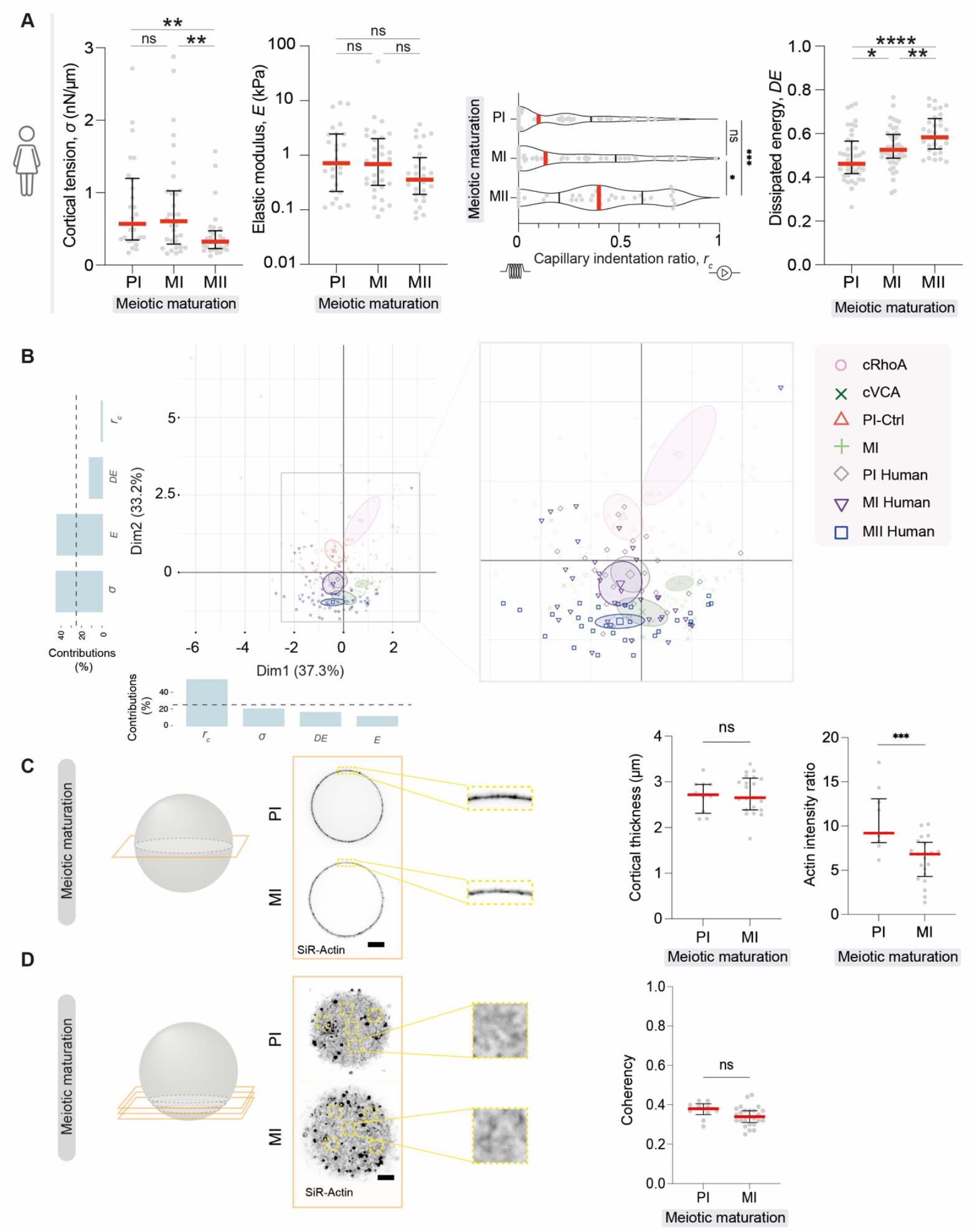
Human and mouse oocyte mechanics diverge, reflecting differences in cortex organization. **A.** Graphs showing the mechanical parameters (cortical tension σ (nN.μm^-1^), elastic modulus E (kPa), capillary indentation ratio r^c^ (a.u), normalized dissipated viscous energy (a.u)) for different stages of human oocyte meiotic maturation (prophase I (PI), meiosis I (MI) and meiosis II (MII)). Data are medians and quartiles with individual data points plotted from four independent experiments. For cortical tension and elastic modulus, 26 PI, 33 MI, and 30 MII were analyzed. For capillary indentation ratio and normalized dissipated viscous energy, 43 PI, 47 MI, and 33 MII were analyzed. The statistical significance of differences between 2 groups was assessed with an unpaired t-test or a nonparametric Mann-Whitney test, depending on whether the data followed a Gaussian distribution. **B.** Principal component analysis of the four mechanical parameters for different stages of mouse oocyte meiotic maturation and engineered mouse oocytes and the different stages of human oocytes meiotic maturation (20 cRhoA, 77 PI-Ctrl, 22 cVCA, 50 MI, 26 human PI, 33 human MI and 30 human MII). Ellipses indicate 95% confidence intervals. The graphs show the percentage contribution of the four mechanical parameters in PCA Dimension 1 (Dim-1) and Dimension 2 (Dim-2). The black dotted line is set at 25%. **C.** Confocal spinning disc images of human oocytes stained with SiR-Actin (black) in prophase I (PI) and meiosis I (MI). One Z plane is shown. Scale bar: 20 μm. Graph showing the cortical thickness and the actin fluorescence ratio between the cortex and the cytoplasm of 10 PI and 21 MI human oocytes. Data are medians and quartiles with individual data points plotted from three independent experiments. Statistical significance of differences between 2 groups was assessed with a parametric t-test. **D.** Confocal spinning disc images of human oocytes stained with SiR-Actin (black) in prophase I (PI) and meiosis I (MI). A projection of six Z planes is shown. Scale bar: 10 μm. Graph showing the coherency of the actin network calculated in 5.33 μm square ROIs of 10 PI and 25 MI oocytes. Data are medians and quartiles with individual data points plotted from three independent experiments. Statistical significance of differences between 2 groups was assessed with a parametric t-test.

The mechanical measurements and the PCA suggested potential differences in the organization of the human oocyte cortex compared to the mouse one. At the prophase I stage, human oocytes appear softer than mouse ones, suggesting that they could have a thicker actin cortex. The characterization of their actin cortex indeed revealed a thicker cortex for human compared to mouse oocytes in prophase I (Figure 3C and Figure 4C). Furthermore, human oocytes in prophase I and meiosis I displayed the same mechanical parameters, suggesting an absence of cortical actin thickening after meiosis I entry in human oocytes, as is normally the case in mouse oocytes^[7]^. Indeed, our results show that human oocytes in prophase I and meiosis I present similar cortex thickness and cortical coherency (Figure 4C, 4D), with an actin density lower in meiosis I compared to prophase I (Figure 4C).

Therefore, as in mouse oocytes, variations in cortex structure correlate with the mechanical properties of human oocytes. However, the evolution of mechanical properties during meiotic maturation differs between human and mouse oocytes, the mechanical profile changing later during meiotic maturation in human compared to mouse oocytes.

## DISCUSSION

In this study, we linked oocyte’s mechanical properties to their actin cortical organization using AFM experiments: a thin actin cortex corresponds to stiff oocytes while a thick actin cortex is associated with softer oocytes. We also described this link between actin cortical organization and mechanical properties in the subfertile case of mouse advanced maternal age and in human oocytes.

We developed an elasto-capillary model to mechanically characterize oocytes probed with a local deformation. This model considers the deformation of a capillary element coupled to an elastic element. While the cell elastic modulus has been extensively investigated at the local scale in numerous AFM studies^[26,27,30]^, capillary deformation has been considered only at the scale of the oocyte’s global deformation^[38]^. Furthermore, previous research suggested that both elastic and capillary response should be considered, especially in cases involving local deformations as in our study^[23,24]^. Indeed, we found that our model better describes the oocyte’s response to a small indentation than models that take into account only one of these non-dissipative responses. Our model could be applied to other cell types as cancer cells, which have so far been mainly characterized using elastic models in AFM^[12]^.

Our AFM measurements resulted in four mechanical parameters from a single oocyte measurement. We combined them using a PCA, to visualize the mechanical phenotypes of each oocyte population studied. This analysis allowed to discriminate between groups that were indistinguishable using a single parameter. For example, cVCA and late meiosis I oocytes have the same cortical tension but appear separate on the PCA. Interestingly, a previous study has combined several mechanical parameters to identify mammalian early embryos with the best developmental potential^[9,39]^. In another study, the clustering and discrimination of metastatic cancer cells was improved by integrating mechanical data with genetic and morphological data^[39]^. Ultimately, mechanical measurements of oocytes could be added to the current multiparametric approach used to identify pathological oocytes in assisted reproductive technology^[40]^.

Focusing on the impact of maternal age, we show that oocytes from old mice are softer than those from young ones with a differently organized actin cortex. A previous study^[41]^ using a different technique showed that oocytes arrested in meiosis II with their *Zona Pellucida* exhibited significantly lower elastic modulus when retrieved from aged mice compared to young ones (3.1 vs. 1.6 kPa respectively), which is in agreement with our results. We show that the cortex of oocytes from aged mice is thicker but less dense than that of oocytes from young mice, suggesting that the cortical actin quantity might be similar or lower in oocytes from aged mice compared to young ones. These results are in agreement with those of a previous study that reported a lower amount of F-actin in aged mice, especially in a sub-cortical region^[42]^. Moreover, coherency analysis shows that cortical actin organization differs significantly between oocytes from young and old mice. These differences could be due to the influence of aging on proteins, including cytoskeletal components^[37]^. Interestingly, it has been shown that cortex softening induces aneuploidy, meaning an abnormal number of chromosomes^[10]^. Therefore, the softening of oocytes associated with maternal age could contribute to the increased aneuploidy rates observed in oocytes from older mammals^[37]^. Investigating the molecular mechanisms underlying these changes in cortex organization would be of great interest.

We probed human oocytes, and show that their mechanical properties evolve differently compared to those of mouse oocytes during meiotic maturation. Cortical tension does not decrease after meiosis I entry in human oocytes, but only in meiosis II. Similarly, their capillary indentation ratio only increases in meiosis II, reflecting a delayed transition between an elastic behavior to a capillary-dominated response. There is a gradual increase in normalized dissipated energy during meiotic maturation, but no significant changes in elastic modulus. One study found that elastic modulus increases between MI and MII in human oocytes, measured by AFM^[26]^. However, these measurements were performed with the *Zona Pellucida*. Another study analyzed the characteristics of the human oocyte cytoplasm through the persistence of the injection funnel after intracytoplasmic sperm injection^[43]^, showing that human oocyte viscosity increases from prophase I to meiosis II, which is consistent with our results. We show that the differences in mechanics measured between mouse and human oocytes are associated with distinct cortical organizations, as highlighted by the absence of cortical actin thickening in human oocytes after entry in meiosis I. In mouse oocytes, the decrease of cortical tension in meiosis I due to cortical actin thickening is essential for the asymmetric in size of the first meiotic division^[7,10,11]^. Human oocytes also divide asymmetrically. However, the lack of cortical actin thickening suggests that alternate mechanisms or cytoskeleton structures may be involved in regulating human oocyte geometry of division.

Finally, this work demonstrates that a thin actin cortex enriched in myosin-II corresponds to stiff oocytes with high cortical tension and elastic modulus. In contrast, a thick actin cortex with less myosin-II is associated with lower cortical tension and elastic modulus. These findings suggest specific links between cortical organization and mechanics in mouse and human oocytes, raising questions about the underlying regulatory mechanisms. *In vitro* studies have shown that actin density and cross-linking control the elasticity of actin networks^[44]^. *In vivo*, altering the orientation of actin filaments, such as transitioning from an actin mesh to fibers, decreases connectivity and affects cell effective elasticity^[45,46]^. Actin dynamics and motor activity control cell tension^[6]^, in agreement with *in vitro* works showing that myosin motors and the organization of actin filaments influence network tension^[47]^. Besides, increasing myosin penetration into the actin cortex results in a thinner cortex and higher cortical tension^[48]^. Therefore, there is generally a subtle crosstalk between actin network structure, dynamics and crosslinking, and motor activity, which we clearly highlight in our experiments. Moreover, some effects could be redundant. For instance, myosin is a motor and a crosslinker, which may explain why tension and elasticity follow the same trend in mouse oocytes. Deciphering these different aspects further would provide a better understanding of how the different mechanical properties of oocytes are regulated and connected.

## MATERIALS AND METHODS

### AFM-indentation measurements

We measured oocytes without their *Zona Pellucida* to study the mechanical properties of the oocyte cortex. Cells were immobilized with an electronic microscopy hexagonal nickel grid (Ref. DT300H-Ni, ∅ 3.05 mm, thickness 18 μm, hole 73 μm; Gilder) attached to the surface of a petri dish filled with medium^[32]^. AFM indentation measurements were performed on a Nanowizard IV AFM (Bruker - JPK Instruments) coupled to a widefield microscope (Zeiss Axio Observer with Hamamatsu sCMOS Flash 4.0 Camera) in contact mode. MLCT-C tips (Bruker, with silicon nitride cantilevers; nominal spring constant = 0.01 N.m^-1^) with a 17° side angle, a 5-μm average height pyramidal tip, and a 20-nm tip radius were used. Before each measurement, the spring constant and the sensitivity of the MLCT tip were calibrated using thermal noise and contact-based methods on the surface of the Petri dishes filled with milli-Q water. Ten force curves were acquired for each oocyte with a 1 μm.s^-1^ approach velocity and a 0.5 nN set point. The measurement takes no more than 1 min for each oocyte.

Force distance curves were converted into force-indentation curves with JPK processing software (version 6.1) by subtracting the cantilever bending from the piezo height to get the accurate vertical tip position, and the contact point was first identified roughly by the “contact point” function. The resulting approach curve was fitted with the elasto-capillary model (Equation (3)), and the contact point was determined precisely by minimizing the Residual Standard Errors (RSE) of the fitting curves. Then, cortical tension *σ* and elastic modulus E, were extracted and used to calculate the capillary indentation ratio parameter *r*_*c*_ (Equation (4)). Finally, the median of cortical tension *σ*, elastic modulus E and capillary indentation ratio *r*_*c*_ was calculated for each oocyte with a minimum of five measurements. The curve fitting and determination of the contact point was done with R studio software (version 2022.12.0+353). The median and the calculation of the quartiles were done with GraphPad Prism 9.5.0 for MacOS, GraphPad Software, La Jolla, CA, USA (version 9.5.0 (525)).

The viscous dissipated energy was calculated by trapezoidal integration of the area between the approach and retraction curves for the positive-indentation region. The value is normalized by the area under the approach curve. All these analyses were done using R studio software (version 2022.12.0+353).

Finally, for each oocyte, the median of normalized viscous energy was calculated with a minimum of five measurements with GraphPad Prism 9.5.0 for MacOS, GraphPad Software, La Jolla, CA, USA (version 9.5.0 (525)).

### Mouse oocyte collection and culture

Ovaries were collected from 11-week-old and 44-56-week-old OF1 female mice. Fully-grown oocytes in prophase I (PI) were extracted by shredding the ovaries in homemade M2 medium^[49]^ supplemented with milrinone (1 μM)^[50]^ or dibutyryl cyclic AMP (dbcAMP)(0.1 mg.mL^-1^)^[51]^ to maintain them in prophase I. The *Zona Pellucida* of PI oocytes was removed by incubating them in a homemade M2 medium supplemented with milrinone (1 μM) or dbcAMP (0.1 mg.mL^-1^) and 0.4% pronase^[52]^. Prophase I exit was triggered by releasing oocytes into a homemade M2 medium. All live culture and imaging were carried out under oil at 37°C.

### Ethical statement

All animal studies were performed in accordance with the guidelines of the European Community and were approved by the French Ministry of Agriculture (authorization D750512).

### Human oocyte collection and culture

The use of human oocytes not arrested in meiosis II and therefore not usable for patients has been approved by the local ethics committee of the Hospices Civils de Lyon (agreement number 22_5725). All patients gave informed consent.

Patients undergoing Assisted Reproductive Technologies (ART) for intracytoplasmic sperm injection (ICSI) had multi-follicular ovarian stimulation. When the ovarian follicles were mature, patients underwent follicular fluid puncture to recover the cumulo-oocyte complexes (COCs). The follicular fluid was screened, and the COCs were cultured in media (Sequential Fert, Origio, Denmark) under oil (Liquid Paraffin, Origio, Denmark) at 37°C, 5% CO2, and 5% O2 in an incubator during 1 hour, and then enzymatically (80UI.mL^-1^ recombinant human hyaluronidase, ICSI Cumulase, Origio) and mechanically denuded. The denuded oocytes were observed under the light microscope to determine their stage. Only oocytes arrested in meiosis II are used for ICSI. The other oocytes in prophase I or meiosis I are unsuitable for ICSI and were used in this study.

The *Zona Pellucida* of human oocytes was removed by incubating them into sequential Fert (Origio, Denmark) with tyrod acid (T1788, Sigma-Aldrich, USA) for 3 seconds. The oocytes were rinsed in sequential Fert and cultured in media (Sequential Fert, Origio, Denmark) under oil (Liquid Paraffin, Origio, Denmark) at 37°C, 5% CO2, and 5% O2 in an incubator.

Meiosis II oocytes were obtained from prophase I or meiosis I oocytes matured *in vitro* for up to 24 hours in a specific medium (MediCult IVM, Origio) supplemented with FSH 75 mUI (Bemfola, GedeonRichter, Hungary), hCG (Ovitrelle, Merck, Germany) and SSS 10% (Fujifilm Irvine Scientific, USA). Oocytes were incubated at 37°C in a controlled atmosphere of 5% CO2 and 5% O2.

### *In vitro* transcription of cRNAs and mouse oocytes microinjection

The following constructs were used: pRN3-EzTD-mCherry-RhoA^[34]^ (cRhoA), gift from Rong Li (Johns Hopkins University, USA), pRN3-EzTD-mCherry^[53]^ (Ctrl) and pRN3-EzTD-mCherry-VCA^[11]^ (cVCA).

Plasmids were linearized using appropriate restriction enzymes. cRNAs were synthesized with the mMessage mMachine kit (Ambion) and subsequently purified using the RNeasy kit (Qiagen) following the manufacturer’s instructions^[54]^. Their concentration was measured using NanoDrop 2000 from ThermoScientific. Their final concentrations were >800 ng.μL^-1^. cRNAs were centrifuged at 4°C for 45 min at 20,000g prior to microinjection into the cytoplasm of oocytes blocked in prophase I in homemade M2 medium supplemented with 1 μM milrinone or 0.1 mg.mL^-1^ dbcAMP at 37°C, using an Eppendorf Femtojet micro-injector^[55]^. After microinjection, cRNA translation was allowed for 4h to allow direct comparison with cortical tension measurements made with micropipette aspiration on oocytes where translation was allowed for 4h^[11]^.

### Fluorescent probes for live imaging

Mouse and human oocytes were incubated for 30 minutes in homemade M2 medium and Fertmedium respectively supplemented with SiR-actin (1 μM) (Spirochrome-SiR-actin Kit (SC001)) and Hoechst (5 ng.mL^-1^) (Sigma-Aldrich H6024) to label F-actin and DNA. For mouse oocytes, immediately after incubation, spinning disk images were acquired using a Plan-APO x40/1.25NA objective on a Leica DMI6000B microscope enclosed in a thermostatic chamber (Life Imaging Service) equipped with a Retiga 3 CCD camera (QImaging) coupled to a Sutter filter wheel (Roper Scientific) and a Yokogawa CSU-X1 Spinning disk. For human oocytes, spinning disk images were acquired using an HCX PL APO x40 and x63 objectives on a Leica DMI4000 microscope with confocal system CSU-W1-T1 Yokogawa enclosed in a thermostatic chamber equipped with a Photometrics PRIME 95B camera. Metamorph software (Universal Imaging) was used to collect data, and ImageJ (NIH) was used to analyze and process data.

### Cortical thickness and fluorescent measurements

Cortical thickness was measured in oocytes incubated with SiR-Actin on a Z-slice going through the widest part of the oocyte to avoid projection artifacts. Thickness was measured by manually placing lines perpendicularly to the cortex in at least four different oocyte cortex regions. The average of these measurements was the cortical thickness of a given oocyte multiplied by the pixel size.

The actin fluorescence ratio was measured manually by measuring the ratio between the mean of the fluorescence signal at the cortex and in the cytoplasm.

### Coherency measurement

Cortical actin network coherency was measured in oocytes incubated with SiR-Actin on the first five cortical z-slices separated by 1 μm and projected together, taking the maximum signal intensity to image the surface of the cortex and have a better view of its organization^[56]^. Coherency was measured in at least 5 square ROIs of 5.33 μm in length and at least 2 square ROIs of 10.6 μm in length using the OrientationJ Analysis plugin in ImageJ which quantifying the spatial correlations of filaments (version 2.14.0/1.54f). The average of these measurements was the cortical coherency of a given oocyte.

### Statistical analysis

The statistical analysis was performed using GraphPad Prism 9.5.0 for MacOS, GraphPad Software, La Jolla, CA, USA (version 9.5.0 (525)). For comparison between two groups, the normality of the values was tested using a D’Agostino-Pearson’s normality test. The statistical significance of differences was assessed by an unpaired t-test (for a normal distribution of the values) or by a nonparametric Mann-Whitney test (for values that did not follow a normal distribution). All tests were performed with a confidence interval of 95%. In all graphs, ns corresponds to not significant, * corresponds to a P-value<0.05, ** to a P-value<0.01, *** to a P-value<0.001 and **** to a P-value<0.0001.

### Principal component analysis

For clustering, we used the median of mechanical parameters. Each point corresponds to the scaled median of all measures for each mechanical parameters per oocyte. The principal component analysis was run with the PCA function from the FactoMineR^[57]^ R package. Associated plots were generated with the factoextra^[58]^ R package. The ellipses correspond to the 95% confidence intervals of the centroid position for each population and non-overlapping 95% confidence ellipses indicate a statistically significant difference between groups.

### Supporting Information

Supporting Information is available Figures S1 and S2.

## Acknowledgments

We thank all members of the Verlhac-Terret and LAMBE labs for discussions, the CIRB animal facility, Tristan Piolot from the Orion facility, Fanny Sibeud for her role in the initiation of this project, and all the members of the LYMIC-PLATIM, especially Simoné Bovio and Elodie Chatre, for all their advices and discussions.

The AFM microscopy was performed at the Orion Platform (member of France–Bioimaging ANR-10-INBS-XX) of the Center for Interdisciplinary Research in Biology (UMR7241/U1050) of the Collège de France. We acknowledge the contribution of SFR Biosciences (Université Claude Bernard Lyon 1, CNRS UAR3444, Inserm US8, ENS de Lyon) PLATIM-LyMIC, especially Simoné Bovio for assistance in AFM experiment and Elodie Chatre with live imaging experiment. This work was supported by DIM ELICIT du Conseil regional d’Ile de France (DIM ELICIT-AAP-2020 to MET), Biomedical Engineering seed grant program (BME to CC), PSL-QLife interdisciplinary program (QLife-CdF-06-2022 to MET) and FRM ECO-Contrat doctoral program (ECO202206015524 to RB). This work has received support from the Fondation Bettencourt Schueller, support under the program « Investissements d’Avenir » launched by the French Government and implemented by the Agence Nationale de la Recherche, with the references: ANR-10-LABX-54 MEMO LIFE, ANR-11-IDEX-0001-02 PSL* Research University.

## Author Contributions

CC and MET conceived the project that was directed by CC, MET and EL. MHV was involved in some aspects of project supervision. RB, LB and YL designed, performed and analyzed all experiments, with the help of GL for AFM. ML built the elasto-capillary model. NT did the PCA. RB wrote the manuscript, which was seen and corrected by all authors.

## Data Availability Statement

All data needed to evaluate the conclusions in the paper are present in the paper and/or the Supplementary Materials. Additional data related to this paper may be requested from the corresponding authors.

## SUPPLEMENTARY FIGURE LEGENDS

**Figure S1.**
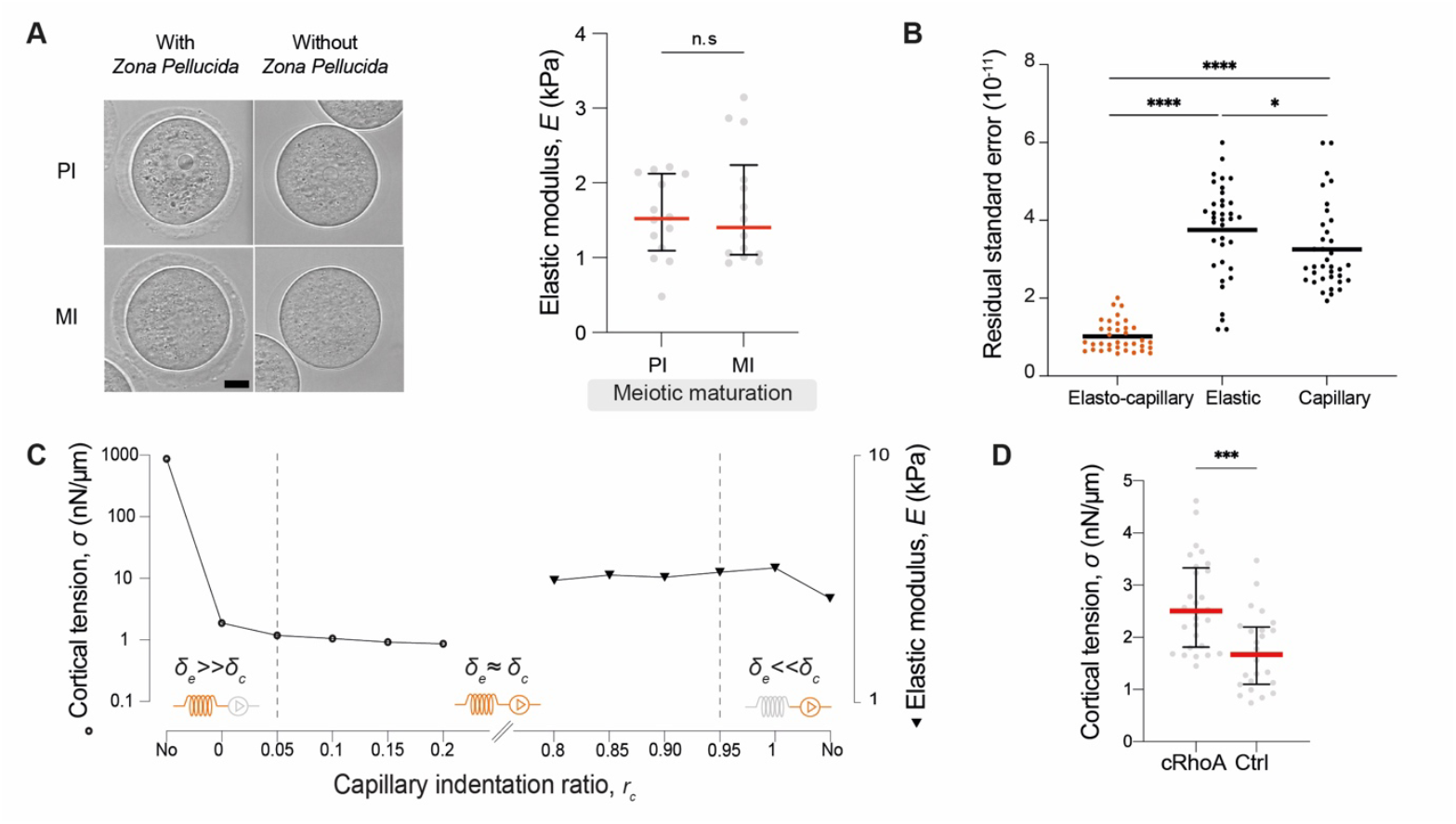
Elastic modulus of mouse oocytes with or without their *Zona Pellucida*, limits and power of the elasto-capillary model, cortical tension of cRhoA oocytes. **A.** Confocal spinning disk transmitted light images of prophase I (PI) and meiosis I (MI) mouse oocytes with (upper panels) and without (lower panels) their *Zona Pellucida*. Scale bar 20 μm. Graph showing the elastic modulus for 14 PI and 14 MI oocytes with their *Zona Pellucida*. Data are medians and quartiles with individual data points plotted from two independent experiments. Statistical significance of differences between 2 groups was assessed with a nonparametric Mann-Whitney test. **B.** Graph plotting the Residual Standard Error (RSE) of elasto-capillary (orange), elastic, and capillary models (both in black). The grey dashed line shows that RSE is reduced with the elasto-capillary model compared to the elastic and capillary models. For each group, 36 oocytes were computed from two experiments (10 PI, 10 early MI, 16 late MI). Data are medians with individual data points plotted. The statistical significance of differences was assessed with a nonparametric Mann-Whitney test. **C.** Graph highlighting the elasto-capillary model limits, showing the mean of cortical tension and elastic modulus as a function of the indentation ratio of the capillary element (r_c_) for oocytes (28 PI, 23 early MI, 27 late MI, 44 MII from at least three experiments). Grey dashed lines show the application limits of the elasto-capillary model (0.05 <r_c_<0.95). **D.** Graph showing the cortical tension measured by micropipette aspiration for 28 cRhoA and 24 Ctrl oocytes. Data are medians and quartiles with individual data points plotted from three independent experiments. Statistical significance of differences between groups was assessed with a parametric t-test.

**Figure S2.**
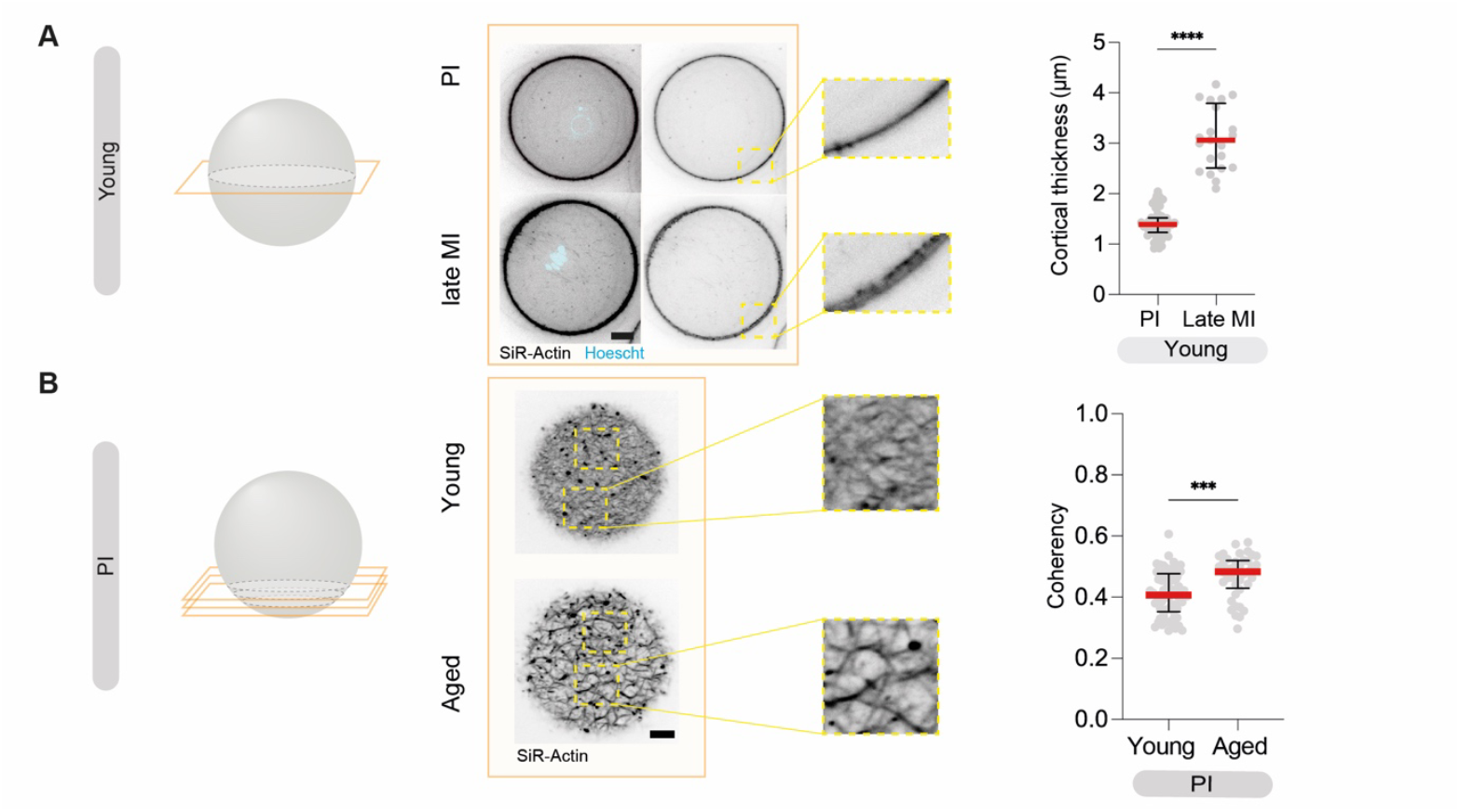
Cortical actin thickness between prophase I and meiosis I mouse oocytes, cortical coherency in oocytes from Young and Aged mice. **A.** Confocal spinning disc images of oocytes in prophase I (PI) and late meiosis I (MI) stained with SiR-Actin (black) and SiR-DNA (blue). Scale bar, 20 μm. One Z plane is shown. Red dotted rectangles highlight magnifications of the oocyte cortex. Graph showing cortical actin thickness for 59 prophase I (PI) and 21 late meiosis I (MI) oocytes. Data are medians and quartiles with individual data points plotted from three independent experiments. Statistical significance of differences between 2 groups was assessed with a parametric t-test. **B**. Confocal spinning disc images of prophase I (PI) oocytes coming from Young (11-week-old) and Aged (44-56-week-old) mice stained with SiR-Actin (black). Scale bar: 5 μm. A projection of five Z planes is shown. Graph showing the coherency of cortex actin network calculated in 10.6 μm square ROIs for 67 prophase I (PI) oocytes coming from Young (11-week-old) and 40 prophase I (PI) oocytes coming from Aged (44-56-week-old) mice. Data are medians and quartiles with individual data points plotted from two independent experiments. Statistical significance of differences between 2 groups was assessed with an unpaired t-test.

## REFERENCES

[1] H. K. Matthews, U. Delabre, J. L. Rohn, J. Guck, P. Kunda, B. Baum, Developmental Cell 2012, 23, 371.

[2] M. Plodinec, M. Loparic, C. A. Monnier, E. C. Obermann, R. Zanetti-Dallenbach, P. Oertle, J. T. Hyotyla, U. Aebi, M. Bentires-Alj, R. Y. H. Lim, C.-A. Schoenenberger, Nature Nanotech 2012, 7, 757.

[3] M. Lekka, K. Pogoda, J. Gostek, O. Klymenko, S. Prauzner-Bechcicki, J. Wiltowska-Zuber, J. Jaczewska, J. Lekki, Z. Stachura, Micron 2012, 43, 1259.

[4] L. Blanchoin, R. Boujemaa-Paterski, C. Sykes, J. Plastino, Physiological Reviews 2014, 94, 235.

[5] E. Evans, A. Yeung, Biophysical Journal 1989, 56, 151.

[6] J.-Y. Tinevez, U. Schulze, G. Salbreux, J. Roensch, J.-F. Joanny, E. Paluch, Proceedings of the National Academy of Sciences 2009, 106, 18581.

[7] A. Chaigne, C. Campillo, N. S. Gov, R. Voituriez, J. Azoury, C. Umaña-Diaz, M. Almonacid, I. Queguiner, P. Nassoy, C. Sykes, M.-H. Verlhac, M.-E. Terret, Nat Cell Biol 2013, 15, 958.

[8] S. M. Larson, H. J. Lee, P. Hung, L. M. Matthews, D. N. Robinson, J. P. Evans, MBoC 2010, 21, 3182.

[9] L. Z. Yanez, J. Han, B. B. Behr, R. A. R. Pera, D. B. Camarillo, Nat Commun 2016, 7, 10809.

[10] I. Bennabi, F. Crozet, E. Nikalayevich, A. Chaigne, G. Letort, M. Manil-Ségalen, C. Campillo, C. Cadart, A. Othmani, R. Attia, A. Genovesio, M.-H. Verlhac, M.-E. Terret, Nat Commun 2020, 11, 1649.

[11] A. Chaigne, C. Campillo, N. S. Gov, R. Voituriez, C. Sykes, M. H. Verlhac, M. E. Terret, Nat Commun 2015, 6, 6027.

[12] M. Liboz, A. Allard, M. Malo, G. Lamour, G. Letort, B. Thiébot, S. Labdi, J. Pelta, C. Campillo, ACS Appl. Mater. Interfaces 2023, 15, 43403.

[13] R. Vargas-Pinto, H. Gong, A. Vahabikashi, M. Johnson, Biophysical Journal 2013, 105, 300.

[14] L. Ramms, G. Fabris, R. Windoffer, N. Schwarz, R. Springer, C. Zhou, J. Lazar, S. Stiefel, N. Hersch, U. Schnakenberg, T. M. Magin, R. E. Leube, R. Merkel, B. Hoffmann, Proc. Natl. Acad. Sci. U.S.A. 2013, 110, 18513.

[15] S. Kasas, G. Longo, G. Dietler, J. Phys. D: Appl. Phys. 2013, 46, 133001.

[16] L. Andolfi, E. Bourkoula, E. Migliorini, A. Palma, A. Pucer, M. Skrap, G. Scoles, A. P. Beltrami, D. Cesselli, M. Lazzarino, PLoS ONE 2014, 9, e112582.

[17] Y. Qi, L. Andolfi, F. Frattini, F. Mayer, M. Lazzarino, J. Hu, Nat Commun 2015, 6, 8512.

[18] K. Haase, A. E. Pelling, J. R. Soc. Interface. 2015, 12, 20140970.

[19] G. Lamour, M. Malo, R. Crépin, J. Pelta, S. Labdi, C. Campillo, ACS Biomater. Sci. Eng. 2024, 10, 1364.

[20] E. Herardot, M. Liboz, G. Lamour, M. Malo, A. Plancheron, W. Habeler, C. Geiger, E. Frank, C. Campillo, C. Monville, K. Ben M’Barek, Stem Cell Rev and Rep 2024, 20, 1340.

[21] F. Rico, P. Roca-Cusachs, N. Gavara, R. Farré, M. Rotger, D. Navajas, Phys. Rev. E 2005, 72, 021914.

[22] R. Garcia, Chem. Soc. Rev. 2020, 49, 5850.

[23] N. Mandriota, C. Friedsam, J. A. Jones-Molina, K. V. Tatem, D. E. Ingber, O. Sahin, Nat. Mater. 2019, 18, 1071.

[24] O. Markova, C. Clanet, J. Husson, Biophysical Journal 2024, 123, 210.

[25] A. Battistella, L. Andolfi, M. Zanetti, S. Dal Zilio, M. Stebel, G. Ricci, M. Lazzarino, Bioengineering & Transla Med 2022, 7, DOI 10.1002/btm2.10294.

[26] L. Andolfi, E. Masiero, E. Giolo, M. Martinelli, S. Luppi, S. dal Zilio, I. Delfino, R. Bortul, M. Zweyer, G. Ricci, M. Lazzarino, Integrative Biology 2016, 8, 886.

[27] J. K. Choi, T. Yue, H. Huang, G. Zhao, M. Zhang, X. He, Cryobiology 2015, 71, 350.

[28] E. Giolo, M. Martinelli, S. Luppi, F. Romano, G. Ricci, M. Lazzarino, L. Andolfi, Eur Biophys J 2019, 48, 585.

[29] T. Shen, Acta Biomaterialia 2019, 10.

[30] L. Andolfi, S. L. M. Greco, D. Tierno, R. Chignola, M. Martinelli, E. Giolo, S. Luppi, I. Delfino, M. Zanetti, A. Battistella, G. Baldini, G. Ricci, M. Lazzarino, Acta Biomaterialia 2019, 94, 505.

[31] A. Chaigne, C. Campillo, R. Voituriez, N. S. Gov, C. Sykes, M.-H. Verlhac, M.-E. Terret, Nat Commun 2016, 7, 10253.

[32] R. Bulteau, L. Barbier, G. Lamour, T. Piolot, E. Labrune, C. Campillo, M.-E. Terret, in Cell Cycle Control (Eds: A. Castro, B. Lacroix), Springer US, New York, NY, 2024, pp. 117–124.

[33] G. G. Bilodeau, Journal of Applied Mechanics 1992, 59, 519.

[34] H. Wang, Y. Li, J. Yang, X. Duan, P. Kalab, S. X. Sun, R. Li, Sci. Adv. 2020, 6, eaaz5004.

[35] D. Cimadomo, G. Fabozzi, A. Vaiarelli, N. Ubaldi, F. M. Ubaldi, L. Rienzi, Front. Endocrinol. 2018, 9, 327.

[36] E. E. Telfer, J. Grosbois, Y. L. Odey, R. Rosario, R. A. Anderson, Physiological Reviews 2023, 103, 2623.

[37] C. Charalambous, A. Webster, M. Schuh, Nat Rev Mol Cell Biol 2023, 24, 27.

[38] E. Fischer-Friedrich, A. A. Hyman, F. Jülicher, D. J. Müller, J. Helenius, Sci Rep 2014, 4, 6213.

[39] K. M. Young, C. Xu, K. Ahkee, R. Mezencev, S. P. Swingle, T. Yu, A. Paikeday, C. Kim, J. F. McDonald, P. Qiu, T. Sulchek, iScience 2023, 26, 106393.

[40] P. Palay, D. Fathi, R. Fathi, Biology of Reproduction 2023, 108, 393.

[41] X. Liu, J. Shi, Z. Zong, K.-T. Wan, Y. Sun, Ann Biomed Eng 2012, 40, 2122.

[42] X. Liu, R. Fernandes, A. Jurisicova, R. F. Casper, Y. Sun, Lab Chip 2010, 10, 2154.

[43] I. Krause, U. Pohler, S. Grosse, O. Shebl, E. Petek, A. Chandra, T. Ebner, Fertility and Sterility 2016, 106, 1101.

[44] M. L. Gardel, J. H. Shin, F. C. MacKintosh, L. Mahadevan, P. Matsudaira, D. A. Weitz, Science 2004, 304, 1301.

[45] F. Eghiaian, A. Rigato, S. Scheuring, Biophysical Journal 2015, 108, 1330.

[46] C. G. Muresan, Z. G. Sun, V. Yadav, A. P. Tabatabai, L. Lanier, J. H. Kim, T. Kim, M. P. Murrell, Nat Commun 2022, 13, 7008.

[47] A.-C. Reymann, R. Boujemaa-Paterski, J.-L. Martiel, C. Guérin, W. Cao, H. F. Chin, E. M. De La Cruz, M. Théry, L. Blanchoin, Science 2012, 336, 1310.

[48] B. A. Truong Quang, R. Peters, D. A. D. Cassani, P. Chugh, A. G. Clark, M. Agnew, G. Charras, E. K. Paluch, Nat Commun 2021, 12, 6511.

[49] M.-H. Verlhac, J. Z. Kubiak, H. J. Clarke, B. Maro, Development 1994, 120, 1017.

[50] A. Reis, H.-Y. Chang, M. Levasseur, K. T. Jones, Nat Cell Biol 2006, 8, 539.

[51] E. Borsuk, J. Z. Kubiak, in Mouse Oocyte Development (Eds: M.-H. Verlhac, M.-E. Terret), Springer New York, New York, NY, 2018, pp. 13–21.

[52] M. Almonacid, in Mouse Oocyte Development (Eds: M.-H. Verlhac, M.-E. Terret), Springer New York, New York, NY, 2018, pp. 145–151.

[53] S. Louvet, J. Aghion, A. Santa-Maria, P. Mangeat, B. Maro, Developmental Biology 1996, 177, 568.

[54] M. E. Terret, C. Lefebvre, A. Djiane, P. Rassinier, J. Moreau, B. Maro, M.-H. Verlhac, Development 2003, 130, 5169.

[55] M.-H. Verlhac, The EMBO Journal 2000, 19, 6065.

[56] E. Nikalayevich, G. Letort, G. De Labbey, E. Todisco, A. Shihabi, H. Turlier, R. Voituriez, M. Yahiatene, X. Pollet-Villard, M. Innocenti, M. Schuh, M.-E. Terret, M.-H. Verlhac, Developmental Cell 2024, 59, 841.

[57] S. Lê, J. Josse, F. Husson, J. Stat. Soft. 2008, 25, DOI 10.18637/jss.v025.i01.

[58] A. Kassambara, F. Mundt, 2016.

